# Cross-cohort analysis reveals conserved gut virome signatures and phage-encoded auxiliary functions in ulcerative colitis

**DOI:** 10.64898/2026.01.21.700755

**Authors:** Hongjae Park, Hyunyoung Jo, Jihye Noh, Yeonjung Lim, Hong Koh, Dong-Woo Lee, IInam Kang, Jang-Cheon Cho

## Abstract

While gut bacteriome dysbiosis is a well-established hallmark of ulcerative colitis (UC), the ecological and functional remodeling of the gut virome conserved across diverse populations remains unclear. Here, by constructing a cross-cohort atlas of the UC-associated fecal virome, we show that viral reorganization closely parallels bacterial dysbiosis, with enrichment of phages targeting disease-associated bacterial taxa. We identified a conserved set of UC-associated viral signatures that robustly distinguished UC from healthy controls using machine-learning-based classification across independent cohorts. Functional profiling further revealed that phages enriched in UC carried higher densities of auxiliary metabolic genes (AMGs) related to virulence and antibiotic resistance than phages depleted in UC, with immune evasion and glycopeptide resistance genes particularly overrepresented. Together, our cross-cohort approach highlights the value of the gut virome-based diagnostic framework for UC and suggests that phage-encoded AMGs may contribute to shaping the gut ecosystem under inflammatory conditions.

## Introduction

Ulcerative colitis (UC) is a major subtype of inflammatory bowel disease (IBD), affecting approximately five million individuals worldwide^1^. UC is characterized by chronic intestinal inflammation resulting from complex interactions among host genetic susceptibility, immune dysregulation, and gut microbiota dysbiosis^2^. Advances in fecal metagenomic sequencing have enabled culture-independent characterization of gut bacterial dysbiosis, including the depletion of short-chain fatty acid (SCFA)–producing bacteria and the enrichment of inflammation-associated, pathogenic, and antibiotic-resistant taxa^3^.

While microbiome research in UC has traditionally focused on bacterial communities, the gut virome has recently gained attention as a critical yet underexplored component of IBD^4^. Growing evidence suggests that alterations in the gut virome are closely linked to bacterial dysbiosis and host immune responses^5^. Notably, experimental transfer of virus-like particles (VLPs) from IBD patients into human microbiota-associated mice has been shown to trigger intestinal inflammation^6^ and exacerbate colitis severity^7^, highlighting the potential involvement of the virome in disease onset and progression. From a diagnostic perspective, the gut virome has also attracted interest as a potential source of disease-associated signals owing to its relative temporal stability in healthy individuals^8,9^. However, viral communities are highly individualized and exhibit substantial variability across geographic and ethnic backgrounds^10^, raising questions about the generalizability of UC-associated virome signatures. Although several studies have reported promising diagnostic performance based on disease-associated virome alterations^11,12^, inconsistent results across independent cohorts underscore the need for cross-cohort approaches that account for population diversity.

Beyond compositional alterations of the gut virome, understanding its functional contribution to UC is crucial for elucidating disease-associated microbial ecology. Given that the vast majority of intestinal viruses are bacteriophages that infect bacteria^13^, their effects on disease-related processes are likely mediated primarily through interactions with their bacterial hosts. One prominent viral strategy involves the expression of auxiliary metabolic genes (AMGs) during infection, which can reprogram host physiology and pathogenicity^14^. Recent studies in Crohn’s disease, another subtype of IBD, have identified phage-encoded AMGs associated with bacterial pathogenicity, such as *pagC* involved in bacterial survival within macrophages, as well as *rfbB* and *rfbC* genes associated with O-antigen diversification and immune evasion^15^. In contrast, the functional contributions of the gut virome in UC, particularly in relation to bacterial dysbiosis and inflammatory environments, remain poorly defined.

To address these gaps, we performed VLP metagenome sequencing to generate targeted gut viral community profiles from fecal samples collected from a South Korean cohort and integrated these datasets with publicly available metagenomes from multiple international cohorts spanning diverse geographic and ethnic backgrounds. This unified cross-cohort dataset enabled the identification of conserved phage–host interaction patterns and UC-associated virome signatures across populations. Leveraging these shared signatures, we establish a noninvasive, virome-based diagnostic framework for UC and uncover phage-encoded auxiliary functions associated with intestinal inflammation.

## Results

### Reconstruction of gut virome profiles across global cohorts

To characterize gut virome configurations associated with UC, we collected fecal samples from 73 South Korean participants, comprising 36 healthy controls and 37 patients diagnosed with UC (Supplementary Table 1). The control group, aged 24–36 years, showed no abnormalities in routine blood tests and had no history of gastrointestinal disorders. UC patients ranged from 16 to 45 years of age, with disease duration spanning from recent onset to 20 years. Fecal samples from all participants were subjected to VLP metagenome sequencing to enable in-depth profiling of the gut virome^16^. To facilitate cross-cohort analyses, we integrated publicly available metagenomic datasets representing diverse geographic and ethnic backgrounds. This comprised seven cohorts across six countries (Fig. 1a and Supplementary Table 2), including VLP metagenomes from Israel (n = 177) and bulk metagenomes from South Korea (n = 105), China (n = 40), the Netherlands (n = 45), Spain (n = 151), and the United States (US, two cohorts; n = 87 and 81, respectively). In total, 759 metagenomes from eight cohorts were combined to form the discovery dataset, which was used to recover viral contigs and identify UC-associated virome signatures. Two additional cohorts from Hong Kong (n = 255) and the United States (n = 140) were reserved as independent datasets for validation analyses.

**Fig. 1.**
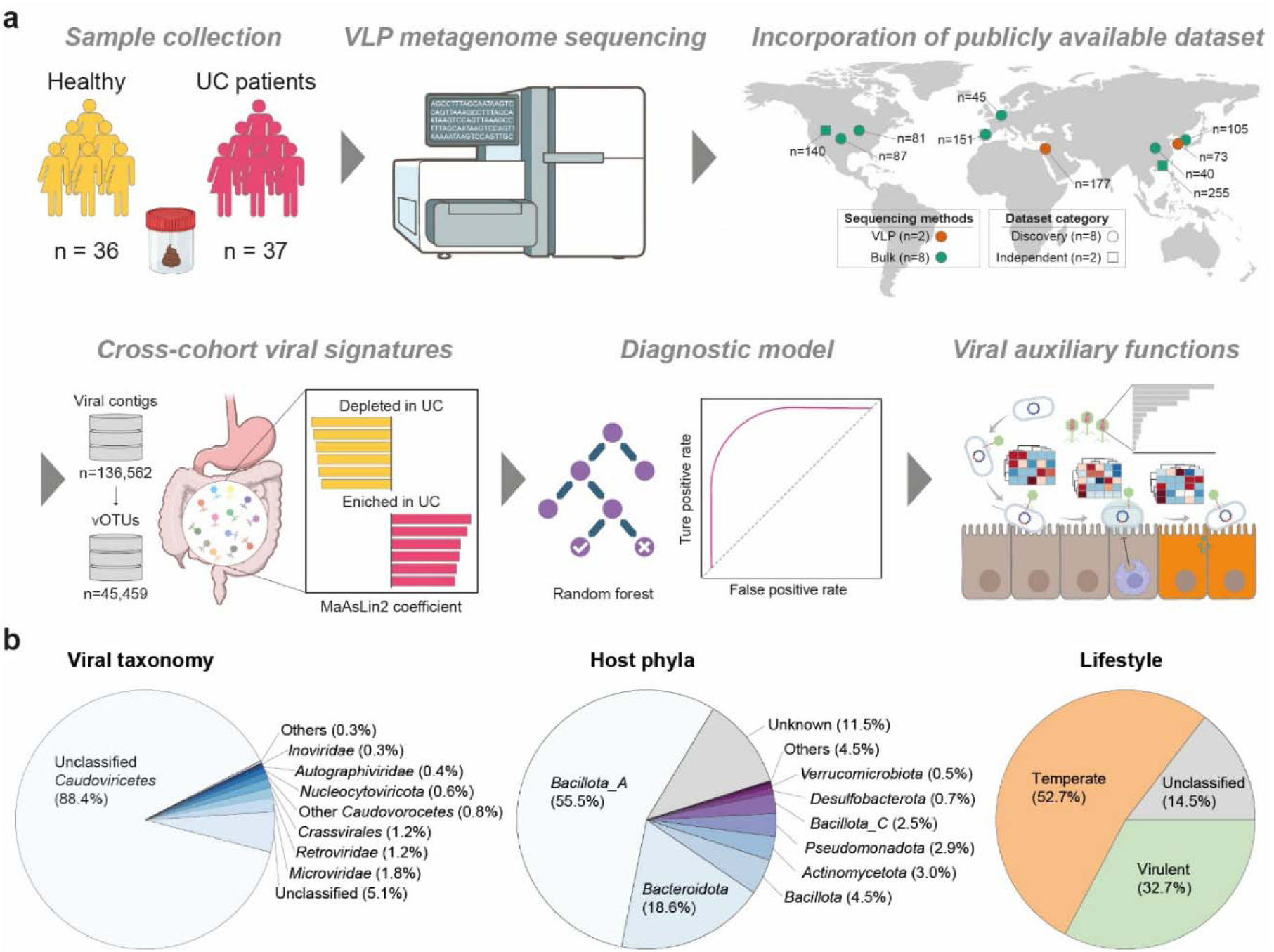
Study design and overview of gut virome reconstruction across multiple cohorts. **a**, Schematic overview of the study workflow. Virus-like particle (VLP) metagenome sequencing was performed on 73 in-house fecal samples from 36 healthy controls and 37 patients with ulcerative colitis (UC). Seven publicly available cohorts were integrated to construct the discovery dataset, with two additional cohorts reserved as an independent validation dataset. Cross-cohort gut virome signatures were identified and used to develop a machine-learning-based diagnostic model. Viral auxiliary metabolic functions associated with disease-related gut inflammation were also characterized. **b**, Summary of identified viral operational taxonomic units (vOTUs) categorized by viral taxonomy (left), predicted bacterial host phylum, (middle), and phage lifestyle (right).

Metagenomic assembly of the discovery dataset yielded 136,562 putative viral contigs (Supplementary Table 3), from which 68,271 viral operational taxonomic units (vOTUs) were defined based on MIUViG criteria^17^ (average nucleotide identity ≥95% and aligned fraction ≥85%). Subsequent quality assessment classified 3.6% of vOTUs as complete and 8.6% as medium to high quality (Extended Data Fig. 1). After excluding vOTUs with undetermined completeness (33.4%), a total of 45,459 high-confidence vOTUs were retained for downstream analyses. Taxonomic classification (Fig. 1b) revealed that the majority of these vOTUs (88.4%) were assigned to unclassified members of the class *Caudoviricetes*, with smaller fractions affiliated with *Microviridae* (1.8%), *Retroviridae* (1.2%), *Crassivirales* (1.2%), and *Nucleocytoviricota* (0.6%). Host prediction was achieved for 83.6% of vOTUs, which predominantly infected bacteria belonging to *Bacillota_A* (55.5%) and *Bacteroidota* (18.6%). Lifestyle analysis further indicated that 52.7% of vOTUs were temperate and 32.7% were virulent (Fig. 1b).

Analysis of vOTU occurrence across samples based on metagenomic read recruitment showed that 16.6% of vOTUs (n = 7,545) were detected across all cohorts, constituting a core set of broadly distributed gut viruses (Extended Data Fig. 2). In contrast, 24.5% (n = 11,151) were detected in only a single cohort, likely reflecting differences in sequencing approaches and depth as well as population-level heterogeneity. Notably, the newly generated South Korean VLP dataset contributed the largest number of cohort-specific vOTUs (n = 3,408).

### Disease-associated remodeling of the gut virome

Non-metric multidimensional scaling (NMDS) of viral community profiles revealed a clear separation between UC patients and healthy controls (PERMANOVA *p*-value < 0.001), indicating pronounced disease-associated compositional shifts in the gut virome (Fig. 2a). Cross-cohort comparisons within the UC and control groups (Fig. 2b) further revealed significant variation in virome composition by geographic origin and sequencing methodologies (VLP vs. bulk metagenomes) (*p*-value < 0.001), indicating substantial cohort-specific effects. These findings emphasize the importance of accounting for cohort-specific heterogeneity when identifying shared viral signatures that are consistently associated with UC. To contextualize these community-level changes within an ecological framework, we first assessed the relative abundances of viral groups according to predicted host phyla (Fig. 2c). Phages infecting *Bacillota_A* and *Bacteroidota* dominated most samples, together accounting for more than 90% of the viral community. These two groups exhibited an inverse relationship (Pearson’s *r* = −0.59, *p*-value < 2.2e−16), with enrichment of *Bacillota_A*-infecting phages generally coinciding with a reduction in *Bacteroidota*-infecting phages (Extended Data Fig. 3). However, the relative proportions of these phyla did not differ markedly between UC and control groups, consistent with previous observations^18^ that phylum-level viral composition largely reflects inter-individual variation rather than robust disease-specific signals.

**Fig. 2.**
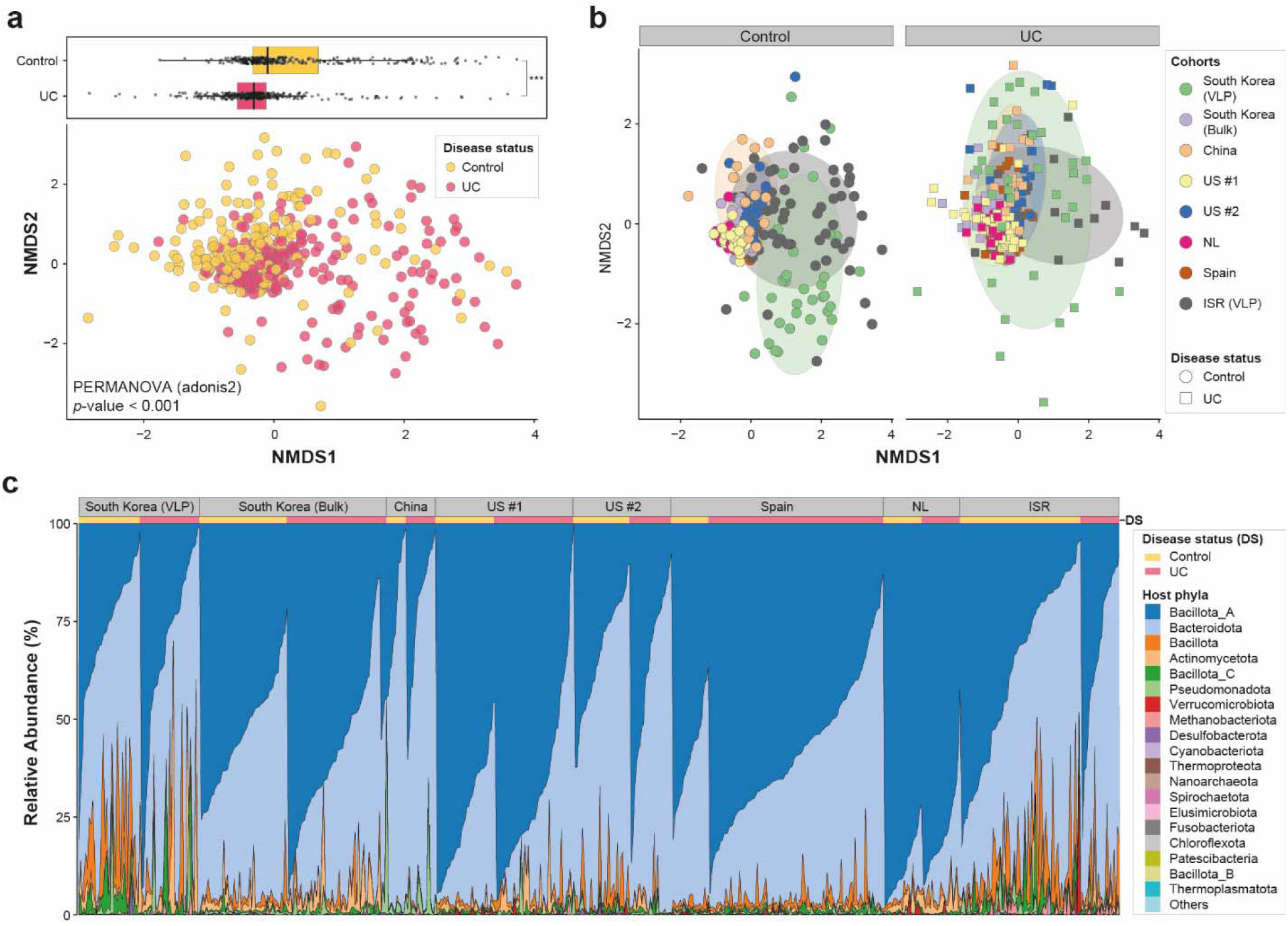
Disease-associated alterations in gut virome community structures. **a**, Non-metric multidimensional scaling (NMDS) ordination of viral community profiles based on vOTU abundance. Boxplots show the distribution of NMDS1 values for healthy controls and UC patients. Wilcoxon test: \*\*\**p* ≤ 0.001. **b**, The same NMDS ordination as in panel a, with samples colored by cohort origin and annotated by disease status. Group-level variation is illustrated with overlaid ellipses representing the 70% confidence region for each cohort. **c**, Stacked bar plot showing viral community composition based on predicted host phyla. Samples are ordered by the relative abundance of phages infecting *Bacillota_A*.

We next applied generalized linear models implemented in MaAsLin2^19^, incorporating viral taxonomy, phage lifestyle, and predicted bacterial host species. This analysis identified significant differences between UC and control groups in the abundances of 14 viral taxa, two phage lifestyle categories, and 348 predicted host bacterial species (151 after excluding taxa without valid species-level annotations) (Fig. 3 and Supplementary Table 4). Notably, the UC group exhibited significant reductions in *Crassvirales*, *Intestiviridae*, *Crevaviridae*, *Microviridae*, and *Mesyanzhinovviridae* (FDR-adjusted *p*-values < 0.05, coefficients < −1.18) (Fig. 3a), several of which, such as *Crassvirales*^20^ and *Microviridae*^21^, have previously been reported to be depleted in IBD. In contrast, *Inoviridae*, *Poxviridae*, *Kyanoviridae*, *Megaviricetes*, *Vilmaviridae*, *Demerecviridae*, *Peduoviridae*, *Autographiviridae*, and unclassified *Caudoviricetes* were significantly enriched in UC (FDR-adjusted *p*-values < 0.05, coefficients > 0.13), consistent with earlier reports linking several of these groups, including *Inoviridae*^22^, *Autographiviridae*^23^, and *Caudoviricetes*^20^, to IBD or other gastrointestinal disorders. Beyond taxonomic differences, UC was associated with marked shifts in phage lifestyles (Fig. 3b). Consistent with previous IBD studies reporting a compositional shift from virulent to temperate phages^11,12^, we observed a significant depletion of virulent phages (FDR-adjusted *p*-value < 0.05, coefficient = −0.19) and a corresponding enrichment of temperate phages (FDR-adjusted *p*-value < 0.05, coefficient = 0.29) in UC.

**Fig. 3.**
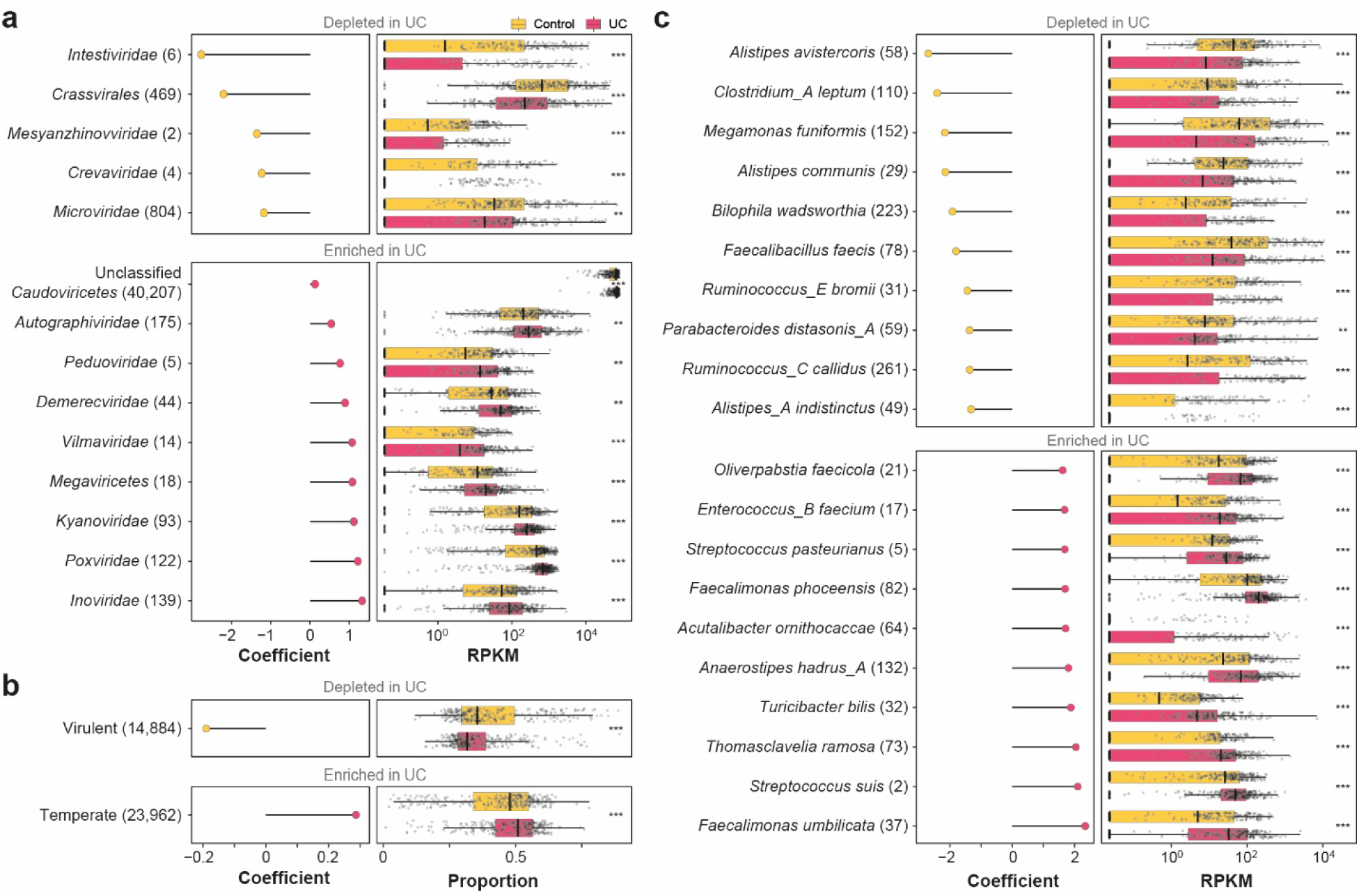
UC-associated alterations in gut virome composition identified by multivariable association analysis. Multivariable association results (coefficients and abundances) obtained using MaAsLin2 general linear model analysis showing significantly altered viral taxonomic groups (**a**), phage lifestyle categories (**b**), and predicted host bacterial species (**c**) between UC patients and healthy controls. For predicted host species, the top 10 UC-depleted and top 10 UC-enriched species with resolved species-level annotations are shown; full results are provided in Supplementary Table 4. The number of vOTUs within each group is indicated in parentheses. Wilcoxon test: \**p* ≤ 0.05, \*\**p* ≤ 0.01, \*\*\**p* ≤ 0.001.

Given the close ecological coupling between phages and their bacterial hosts, alterations in the gut virome often parallel changes in the bacteriome^12^. Consistently, UC samples showed significant depletion of phages predicted to infect 114 bacterial species (FDR-adjusted *p*-values < 0.05, coefficients < −0.25), alongside enrichment of phages targeting 234 distinct species (FDR-adjusted *p*-values < 0.05, coefficients > 0.25) (Fig. 3c and Supplementary Table 4). Phages depleted in UC were primarily associated with fiber-degrading and SCFA-producing commensals, including multiple species of *Alistipes*^24^, *Parabacteroides*^25^, and *Gemmiger*^26^, as well as lactic acid bacteria (*Sharpea*^27^*, Ligilactobacillus*^28^, and *Lactococcus*^29^) and *Akkermansia muciniphila*^30^, a mucin-degrading bacterium associated with epithelial barrier integrity and host metabolic regulation. In contrast, phages enriched in UC were frequently associated with opportunistic and inflammation-linked taxa, including *Streptococcus*^31^, *Enterocloster*^32^, and *Prevotella*^33^, as well as well-established pathobionts such as *Clostridioides difficile*^34^ and the ESKAPE pathogen *Enterococcus faecium*^35^. Although *Blautia* species are commonly considered commensals^36^, phages predicted to infect 27 of 29 different *Blautia* species were enriched in UC (Supplementary Table 4), suggesting a context-dependent association consistent with previous reports implicating *Blautia obeum* in IBD^37^. Collectively, these results indicate that UC-associated virome remodeling parallels well-characterized bacterial dysbiosis, marked by the depletion of health-associated commensals and the expansion of opportunistic taxa linked to mucosal inflammation.

### Cross-cohort virome signatures improve noninvasive UC diagnosis

To resolve disease-associated viral signals at higher resolution, we extended the generalized linear model analysis to individual vOTUs and identified 1,698 vOTUs as robust virome signatures distinguishing UC from healthy controls (Supplementary Table 5). Of these, 1,027 vOTUs were enriched and 671 were depleted in UC (Fig. 4a). To assess their diagnostic potential, 80% of the discovery dataset was randomly assigned to a training set, with the remaining 20% reserved for a validation set. A machine learning (ML)-based model built on the training set using five-fold cross-validation (*see* Methods for details) demonstrated that the identified virome signatures distinguished UC from controls with high accuracy (AUC = 0.865; sensitivity = 80.8%; specificity = 75.9%; Fig. 4b) and retained strong performance in the validation set (AUC = 0.892; sensitivity = 93.1%; specificity = 60.0%; Fig. 4c). Importantly, classifier performance was consistent across populations from distinct geographic regions, achieving AUCs of 0.943 in Asian cohorts and 0.818 in Western cohorts (Fig. 4c), supporting the cross-cohort robustness of these virome signatures. Further evaluation using two independent datasets from the United States and Hong Kong confirmed the generalizability of the classifier, achieving AUCs of 0.855 (Fig. 4d) and 0.907 (Fig. 4e), respectively.

**Fig. 4.**
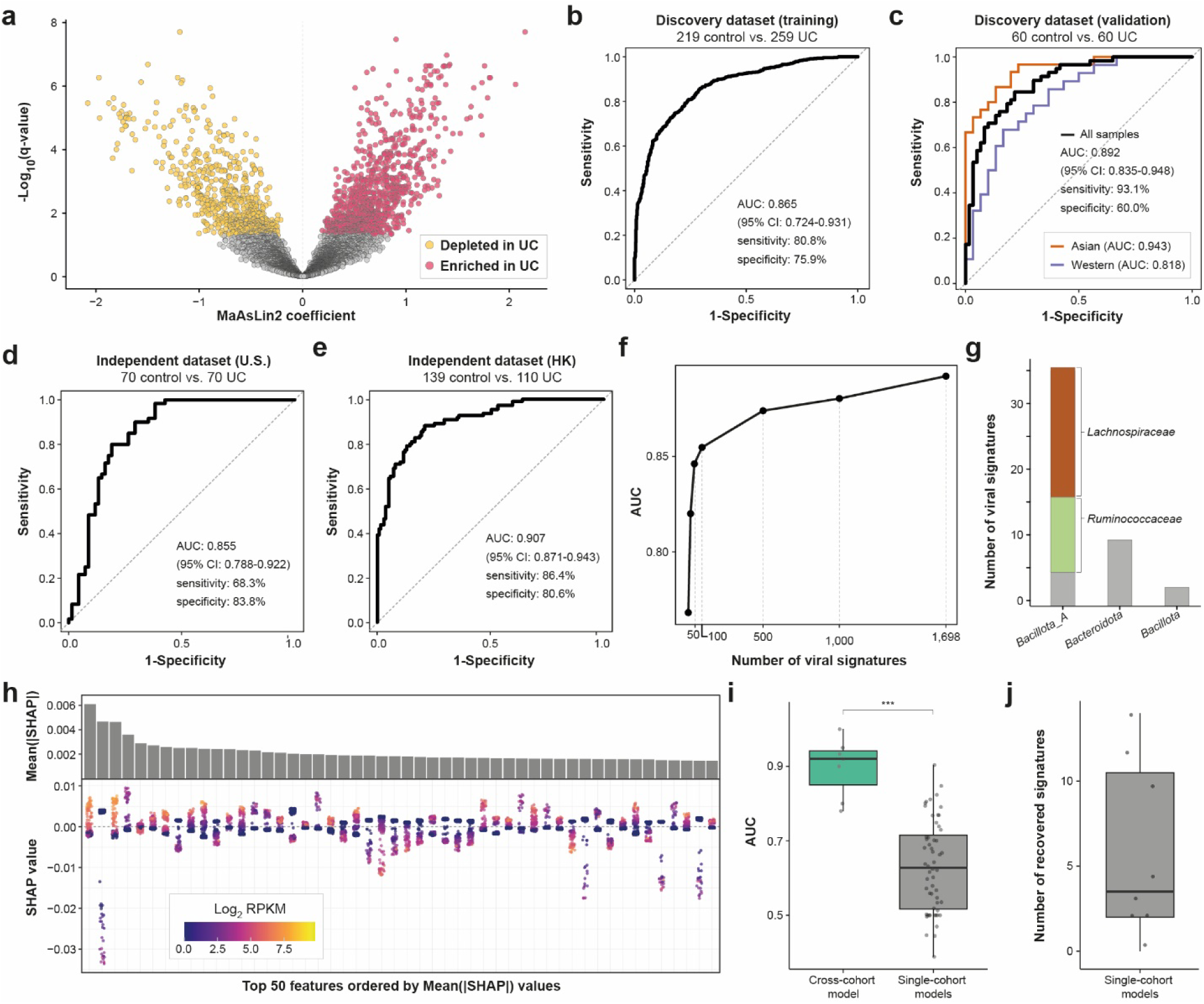
Cross-cohort identification of gut virome signatures and development of an ML–based diagnostic model for UC. **a**, Volcano plot showing shared UC-associated virome signatures identified across cohorts using MaAsLin2. Each point represents a vOTU. **b**, Receiver operating characteristic (ROC) curve showing classifier performance in the training set of the discovery dataset. **c**, ROC curves for the validation set of the discovery dataset, shown for all samples and stratified by geographic region (Asian and Western cohorts). **d-e**, ROC curves showing classifier performance in independent datasets from the United States (d) and Hong Kong (e). **f**, Classification performance (AUC) in the validation set as a function of the number of virome signatures tested in the model. **g**, Predicted bacterial host phyla for the top 50 virome signatures. **h**, SHAP value plot showing the contribution of the top 50 virome signatures across samples. Each point represents a sample and is colored by the log_2_-transformed RPKM value of the corresponding viral signature. **i**, Comparison of diagnostic performance (AUC) between the cross-cohort model and single-cohort models, evaluated across individual cohorts. Wilcoxon test: \*\*\**p* ≤ 0.001. **j**, Number of top 50 cross-cohort signatures recovered by single-cohort models.

To determine the minimal set of viral features required for robust classification, we ranked vOTUs by mean Shapley additive explanations (SHAP) values (Supplementary Table 6), and the least informative signatures were iteratively removed. Classification performance in the validation set declined sharply when fewer than the top 50 vOTUs were retained (Fig. 4f), indicating that this subset represents the lower bound for robust model performance. Host prediction analysis of these top 50 signatures revealed that most were predicted to infect *Bacillota_A* (n = 36), followed by *Bacteroidota* (n = 9) and *Bacillota* (n = 2) (Fig. 4g). At finer taxonomic resolution, 32 of the 50 signatures were associated with the family *Lachnospiraceae* (n = 20) or *Ruminococcaceae* (n = 12). A SHAP-based feature contribution plot further illustrated the direction and magnitude of each vOTU’s contribution to UC classification (Fig. 4h). Among the top 50 signatures, two vOTUs were affiliated with *Nucleocytoviricota* (Supplementary Table 6), although detailed characterization was limited by incomplete genome assemblies (<9 kb), likely reflecting constraints of short-read sequencing.

To directly assess the advantage of a cross-cohort framework over single-cohort models, we re-identified disease-associated virome signatures within each cohort independently, trained cohort-specific classifiers using these signatures, and evaluated their performance across other cohorts. All single-cohort models exhibited substantially reduced performance relative to the cross-cohort model, with a median AUC of 0.628 (Fig. 4i). This reduction was primarily attributable to limited recovery of the top 50 signatures defined by the cross-cohort model, with single-cohort models recapturing an average of only 3.5 of these features (Fig. 4j). Although 43 of the 50 signatures were recovered in at least one single-cohort model (Supplementary Table 7), the remaining seven signatures were not detected in any single-cohort model, indicating that a subset of UC-associated virome signals becomes apparent only through cross-cohort integration.

### Phages modulate bacterial fitness under inflammatory conditions

Despite marked community-level remodeling of the gut virome observed in UC, the functional consequences accompanying these changes remain incompletely characterized. Given the expansion of temperate phages in UC, which are known to carry broader repertoires of AMGs than virulent phages^38^, we examined whether UC-enriched phages exhibit distinct functional profiles. Because AMGs encompass diverse functional categories, subsequent analyses focused on virulence factor genes (VFGs) and antimicrobial resistance genes (ARGs), which are most directly linked to bacterial fitness under host inflammatory conditions^39^. To improve functional annotation of viral genes beyond the limitations imposed by low sequence similarity, we employed a protein structure-guided annotation framework, Phold^40^, which integrates the ProstT5 protein language model^41^ with structural alignment using Foldseek^42^. This approach enabled functional annotation of 38% of the 1.6 million viral genes, more than doubling the annotation yield achieved using the sequence homology-based method implemented in Pharokka^42^ (Supplementary Table 8). Comparative analyses of the gene repertoires of vOTUs enriched (n = 1,027) and depleted (n = 671) in UC revealed a significant overrepresentation of both VFGs and ARGs among UC-enriched phages (Fisher’s exact test *p*-value < 0.001). To account for variation in genome completeness in our dataset, we calculated contig length-normalized gene density (AMGs per Mb), revealing that UC-enriched phages carried 2.4-fold higher densities of VFGs and 3.4-fold higher densities of ARGs than UC-depleted phages (Fig. 5a).

**Fig. 5.**
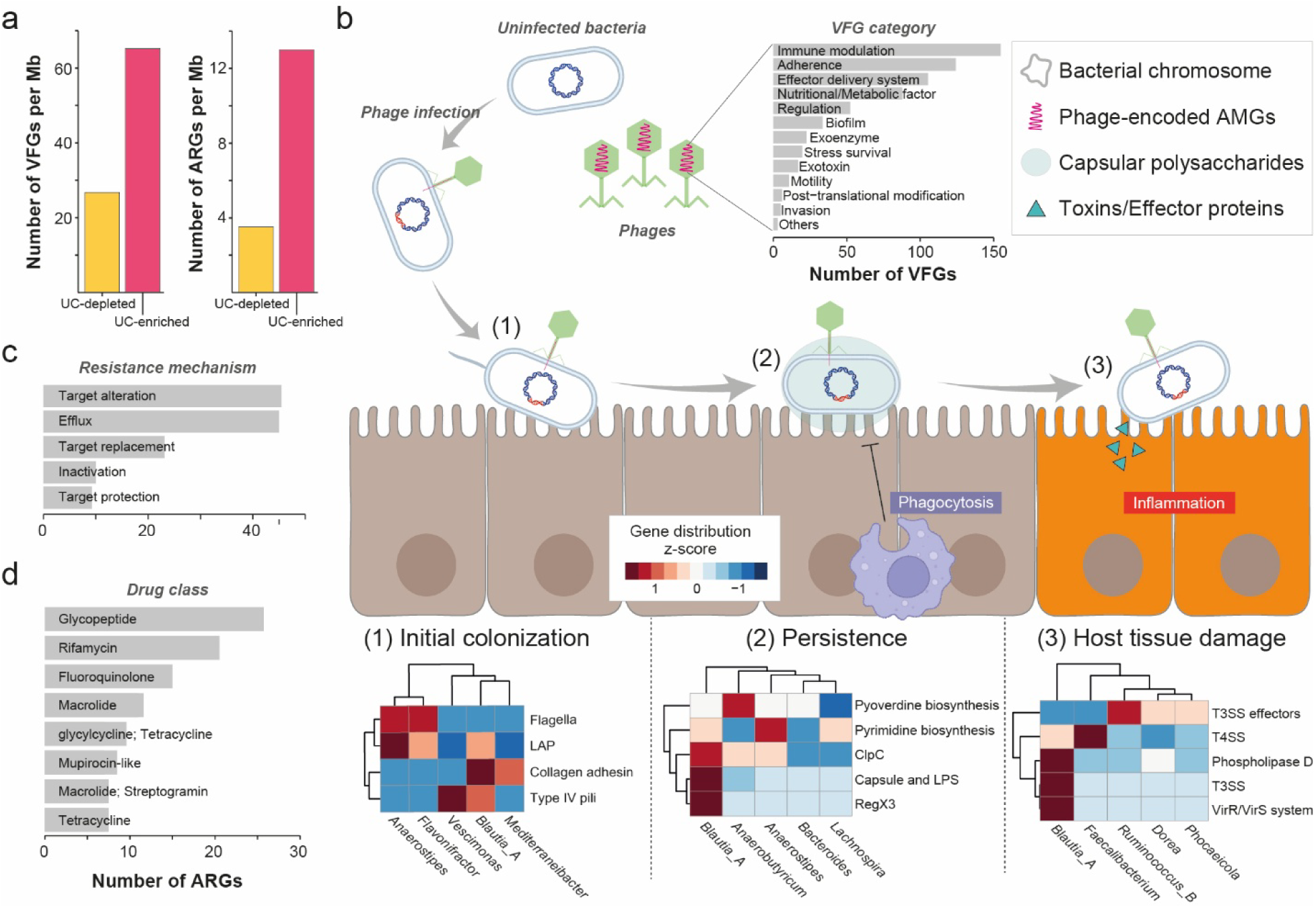
Phage-encoded auxiliary metabolic genes (AMGs) associated with bacterial fitness in UC. **a,** Contig length–normalized densities of virulence factor genes (VFGs) and antimicrobial resistance genes (ARGs) in UC-enriched and UC-depleted phages, expressed as AMGs per Mb. **b,** Functional distribution of VFGs encoded by UC-enriched phages. VFGs are grouped according to major functional categories and mapped onto three stages of host interaction: initial colonization, long-term persistence, and host tissue damage. The heatmap displays the most abundant VFG subcategories and their associated predicted bacterial hosts. **c,** Distribution of ARGs encoded by UC-enriched phages across resistance mechanisms. **d,** Distribution of ARGs encoded by UC-enriched phages across antibiotic drug classes.

We next assessed whether these quantitative differences reflected qualitative shifts in virulence-related functional repertoires. The distribution of functional categories of VFGs was largely comparable between UC-enriched (Fig. 5b) and UC-depleted phages (Extended Data Fig. 4), indicating that UC-associated viromes are primarily characterized by an increased load of virulence-related functions rather than the emergence of novel VFG classes. In UC-enriched phages, VFGs spanned functional categories corresponding to major stages of host interaction^43^, including initial colonization, long-term persistence, and host tissue damage (Fig. 5b and Supplementary Table 9). Genes associated with initial colonization included adherence factors such as the collagen adhesin *cna*^44^ and *lap*^45^, as well as motility-related determinants including pili and flagellar genes. VFGs linked to long-term persistence comprised genes involved in immune evasion, nutrient acquisition (pyoverdine^46^ and pyrimidine biosynthesis), and stress tolerance (*clpC*^47^ and *regX3*^48^). Notably, genes involved in capsule biosynthesis and lipopolysaccharide (LPS) modification (*rfb*^49^*, rff*^50^, and *rml*^51^ clusters) were strongly overrepresented (Extended Data Fig. 5), consistent with functions associated with masking bacterial surface antigens and evading phagocytosis. Host damage-associated functions included effector delivery systems, including type III (T3SS) and type IV secretion systems (T4SS), as well as exotoxins such as hemolysin^52^ and phospholipase D^53^. Among these, T4SS components were particularly enriched (Extended Data Fig. 5), with *virD4*, a coupling protein critical for effector translocation^54^, emerging as the most frequently detected component (Supplementary Table 9).

Consistent with the pattern observed for VFGs, ARG repertoires also exhibited quantitative rather than qualitative shifts (Fig. 5c-d and Extended Data Fig. 6). UC-enriched phages exhibited higher overall ARG densities while maintaining functional category distributions comparable to those of UC-depleted phages. Among ARG functional categories (Fig. 5c and Supplementary Table 10), antibiotic target alteration and efflux mechanisms^55^ were most prevalent. When stratified by drug class (Fig. 5d), genes associated with glycopeptide (*vanH*, *vanR*, and *vanS*^56^) and rifamycin resistance (*rpoB2*^57^) were among the most abundant ARGs, both of which correspond to antibiotic classes commonly used in IBD management^58^, followed by fluoroquinolone resistance determinants (*cdeA*^59^). ARGs conferring resistance to macrolides, glycylcyclines, tetracyclines, and mupirocin were also detected at lower frequencies.

We next investigated which bacterial taxa were most frequently associated with the acquisition of these phage-encoded AMGs. Phages predicted to infect *Blautia* and *Mediterraneibacter* most frequently carried genes encoding collagen adhesins linked to colonization, whereas phages targeting *Anaerobutyricum* were distinguished by preferential presence of genes involved in the biosynthesis of the iron-scavenging siderophore pyoverdine (Fib. 5b). In addition, phages infecting *Blautia* were enriched for genes encoding phospholipase D, T3SS components, and the VirR/VirS^60^ regulatory system. Distinct genus-specific patterns were also observed for ARGs (Extended Data Fig. 7). Phage-borne glycopeptide resistance genes were broadly distributed across multiple genera, including *Blautia_A*, *Lachnospira*, *Anaerostipes*, and *Ruminococcus_B*. Rifamycin resistance genes were preferentially associated with *Blautia_A*, whereas fluoroquinolone resistance was most strongly linked to *Blautia_A* and *Bariatricus*. In contrast, *Phocaeicola*, *Streptococcus*, and *Agathobacter* showed strong associations with tetracycline resistance genes. These non-random distributions of AMGs indicate that phages preferentially equip specific bacterial taxa with specialized functional modules.

Finally, genome-resolved analysis of representative complete vOTUs enriched in UC provided insight into the organization of core and accessory gene repertoires within individual phage genomes. The two largest vOTUs, UC-082_C12519 (91 kb) and SRR6468689_C7150 (41 kb), predicted to infect *Bacteroides thetaiotaomicron* and *Blautia_A obeum*, respectively, exhibited markedly expanded accessory gene complements. Both genomes carried *parA*, *parB*, and *pcfF* genes associated with plasmid-like maintenance^61,62^, reverse transcriptases associated with diversity-generating retroelements^63^, and Paratox proteins implicated in host quorum sensing interferenc^64^. UC-082_C12519 displayed an unusual DNA recombination architecture, encoding at least four tyrosine recombinases alongside a DDE-type integrase^65^. The coexistence of multiple integrase types together with plasmid-like maintenance genes suggests flexibility in intracellular persistence strategies. In contrast, SRR6468689_C7150 encoded a distinct repertoire of host-interactive and stress-response functions, including a hyaluronidase linked to extracellular matrix degradation and inflammatory responses^66^, phosphoadenosine phosphosulfate (PAPS) reductase involved in sulfur metabolism^67^, the RNA polymerase sigma factor SigW linked to stress responses^68^, and Pf4r, a protein associated with superinfection exclusion^69^. Together, these genome-level features highlight the diversity of auxiliary gene repertoires encoded by UC-associated phages with potential relevance for conferring a competitive advantage to host bacteria.

## Discussion

In this study, we constructed a cross-cohort atlas of the UC-associated gut virome by integrating VLP metagenomes with seven publicly available global datasets (Fig. 1). The recovery of thousands of vOTUs unique to the VLP dataset highlights the persistent underrepresentation of viral diversity in bulk metagenomic studies and demonstrates the value of targeted virome sequencing for resolving low-abundance and previously uncharacterized phages (Extended Data Fig. 2). By unifying datasets across sequencing methodologies and populations, our analysis provides one of the most comprehensive and globally representative virome resources for UC to date, enabling the identification of reproducible disease-associated viral signals that extend beyond cohort-specific effects (Figs. 3 and 4).

A central finding of this work is that host-informed classification substantially improves the interpretability of virome alterations in UC. Although the majority (88.4%) of vOTUs in our catalog were classified as members of unclassified *Caudoviricetes* (Fig. 1b), limiting the resolution of taxonomy-based enrichment signals, incorporating phage–host associations revealed disease-associated patterns that closely mirrored well-characterized bacteriome dysbiosis^3^ (Fig. 3). Specifically, UC was characterized by a depletion of phages predicted to infect health-associated commensals and a concomitant enrichment of phages targeting opportunistic and inflammation-associated bacterial taxa. These coordinated shifts suggest that virome remodeling in UC is tightly coupled to bacterial community restructuring and is consistent with ecological dynamics such as the Piggyback-the-Winner regime^38^, in which temperate phages preferentially proliferate alongside expanding bacterial hosts under disturbed conditions. Together, these findings emphasize the importance of host-resolved approaches for interpreting virome dynamics in inflammatory diseases.

Building on these ecological insights, integrating multiple cohorts markedly improved the robustness and generalizability of virome-based disease classifiers (Fig. 4). Diagnostic models trained on single cohorts exhibited inconsistent performance when applied across populations, reflecting pronounced cohort-specific variation in viral community composition. In contrast, a cross-cohort framework enabled the identification of shared UC-associated vOTUs that collectively achieved strong diagnostic performance across diverse cohorts. These findings reinforce the notion that reliable microbiome-based diagnostic signatures should be derived from heterogeneous datasets to disentangle disease-relevant signals from geographic, technical, and lifestyle-associated confounders^70^. Although our classifier performed well in both Asian and Western cohorts, extending this framework to additional underrepresented populations will be necessary to further refine model robustness and translational applicability.

Beyond diagnostic potential, our results provide functional insight into how phages may modulate bacterial fitness in the UC gut environment. By applying a structure-guided annotation strategy, we substantially increased the fraction of viral genes with predicted functions, enabling systematic comparisons of AMG content between UC-enriched and UC-depleted phages (Fig. 5). UC-enriched phages carried significantly higher densities of both VFGs and ARGs, with differences driven predominantly by quantitative increases rather than the emergence of novel functional categories. These findings indicate that UC-associated virome remodeling is characterized by an increased functional load, with phages serving as reservoirs of adaptive traits for host bacteria associated with UC. Notably, AMGs were non-randomly distributed across predicted bacterial hosts, suggesting that phages preferentially associate specific functional modules with particular bacterial taxa rather than functioning as indiscriminate gene shuttles, thereby reinforcing niche-specific adaptive strategies. For example, phages predicted to infect *Blautia*, *Mediterraneibacter*, and *Anaerobutyricum* were preferentially associated with distinct functional modules linked to colonization, nutrient acquisition, and immune interaction (Fig. 5b). Building on these observations, genome-resolved analyses of representative UC-enriched phages further revealed diverse auxiliary functions, including Paratox, hyaluronidase, PAPS reductase, and the stress-response regulator SigW (Fig. 6), collectively supporting a potential role for phages in modulating host-associated bacterial functions in the inflamed gut environment.

**Fig. 6.**
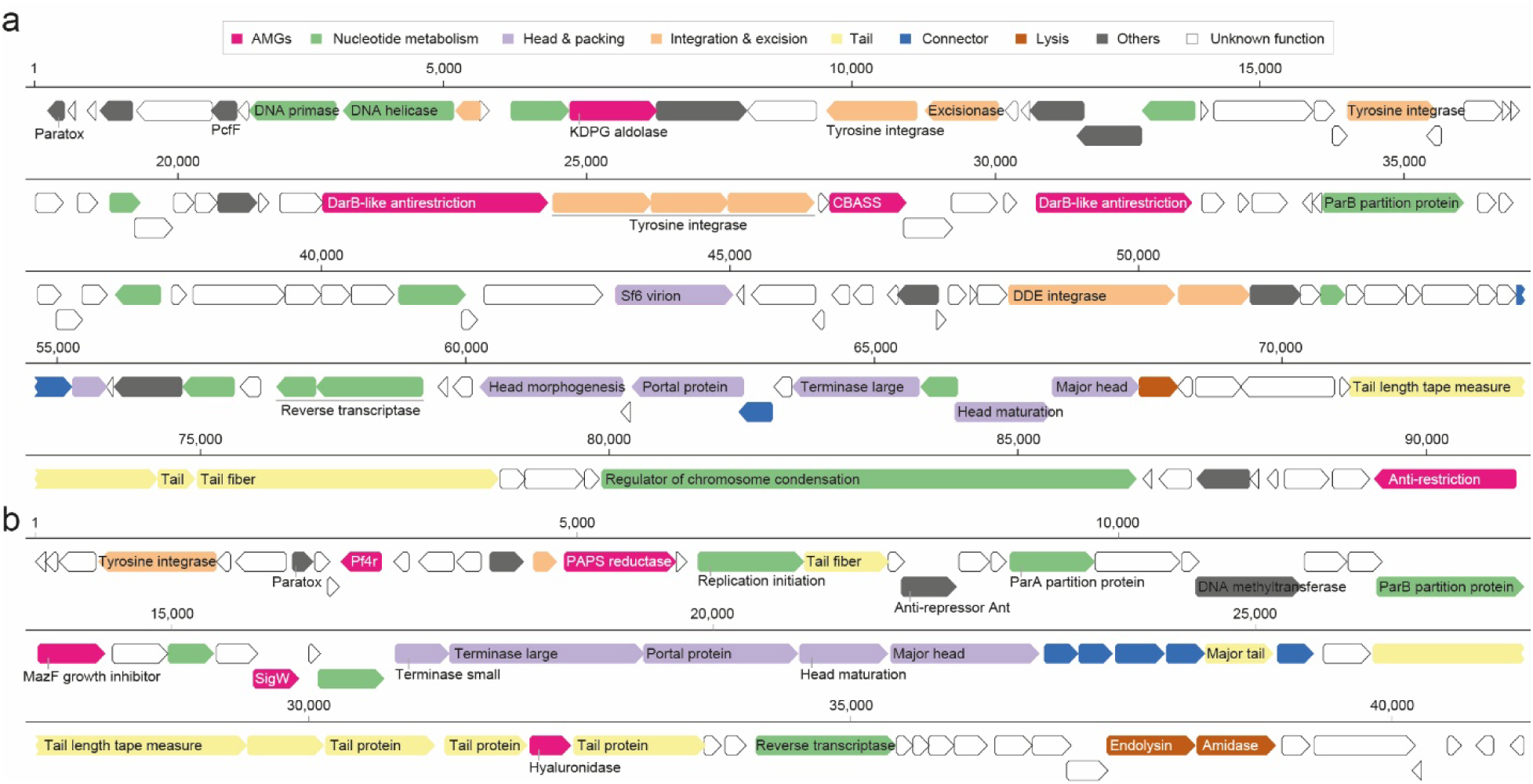
Genome organization of auxiliary gene repertoires in representative complete phage genomes enriched in UC. **a**, Genome map of UC-082_C12519, predicted to infect *Bacteroides thetaiotaomicron*, comprising 132 predicted genes. **b**, Genome map of SRR6468689_C7150, predicted to infect *Blautia_A obeum*, comprising 73 predicted genes. Genes are color-coded by major functional categories, with genes of unknown functions shown in white. Functional annotation was performed using Phold and further refined with HHpred.

Despite these advances, several limitations should be acknowledged. First, the incompleteness of many viral assemblies constrains the reconstruction of full gene neighborhoods and functional modules, likely leading to an underestimation of AMG diversity. Long-read sequencing and improved assembly strategies will be critical for resolving complete phage genomes at scale^71^. Second, functional predictions remain inference-based, and experimental validation will be required to confirm the activity and ecological consequences of phage-encoded AMGs *in vivo*. Finally, although our diagnostic framework achieved robust binary classification (UC versus healthy controls), it was not designed to resolve disease severity or predict disease progression. Incorporating longitudinal sampling and detailed clinical metadata will therefore be essential for evaluating the utility of virome-based signatures in disease stratification, prognosis, and treatment monitoring.

## Methods

### Subject recruitment and sample collection

Stool samples were obtained from healthy South Korean individuals (n = 36) and patients with UC (n = 35), aged 16–45 years. Participants were recruited following approval by the Institutional Review Board (IRB) of Severance Hospital, Yonsei University Health System (IRB No. 4-2020-1487). Written informed consent was obtained from all participants before enrollment. Each participant completed a questionnaire collecting demographic information, medical history, current medication use, and gastrointestinal symptoms potentially influencing gut microbiota composition. Participants who were newly diagnosed with UC during the study period, those with major underlying diseases (e.g., malignancy, multiorgan failure, or peptic ulcer), pregnancy, or abnormal blood or stool screening results were excluded. Additional exclusion criteria included the use of medications known to affect the gastrointestinal tract that could not be discontinued for at least seven days before sampling (e.g., proton pump inhibitors, antacids, antibiotics). Stool samples were collected in sterile conical tubes, transported to the Yonsei Severance Fecal Microbiota Transplantation (FMT) Center (Seoul, South Korea), and stored at −80 °C until further processing.

### VLP enrichment, DNA extraction, and metagenomic sequencing

Approximately 3 g of stool was homogenized in 30 mL of SM buffer (50 mM Tris-HCl, 100 mM NaCl, 8 mM MgSO, pH 7.5) by vortexing for 5 min. The homogenate was centrifuged at 5,000 × *g* for 10 min at 4 °C to remove large particles, and this process was repeated twice. The resulting supernatant was filtered through 0.45 µm PVDF syringe filters (Millipore, Darmstadt, Germany) to remove residual cells and cellular debris. VLPs were precipitated by adding NaCl to a final concentration of 1 M and polyethylene glycol 8000 (PEG 8000; Sigma-Aldrich, St. Louis, MO, USA) to a final concentration of 10% (w/v). The mixture was incubated at 4 °C for 16 h and subsequently centrifuged at 10,000 × g for 20 min at 4 °C. The viral pellet was resuspended in 1 mL of SM buffer and extracted with an equal volume of chloroform (Sigma-Aldrich, St. Louis, MO, USA). The mixture was vortexed briefly and centrifuged at 1,000 g for 10 min at 4 °C to remove remaining bacterial or host-derived contaminants. The aqueous phase was then treated with DNase I (10 U mL^-1^) and RNase A (10 U mL^-1^) (Thermo Fisher Scientific, Waltham, MA, USA) at 37 °C for 45 min to degrade residual bacterial and host nucleic acids.

Enzymes were subsequently heat-inactivated at 70 °C for 10 min. Viral DNA was extracted using the DNeasy Blood & Tissue Kit (Qiagen, Hilden, Germany) following the manufacturer’s protocol. DNA yield and purity were assessed using a Qubit fluorometer (Thermo Fisher Scientific, Waltham, MA, USA). Sequencing libraries were prepared using the NEBNext Ultra II FS DNA Library Prep Kit (New England Biolabs, Ipswich, MA, USA) and sequenced on the Illumina NovaSeq 6000 and NextSeq 2000 platforms at CJ Bioscience (Seoul, South Korea).

### Collection of public metagenomic datasets

Publicly available metagenomic datasets were obtained from the NCBI Sequence Read Archive (SRA; https://www.ncbi.nlm.nih.gov/sra) and GMrepo (https://gmrepo.humangut.info). Studies lacking raw sequencing data or explicit case–control annotations were excluded. The final dataset comprised seven cohorts in the discovery set, including metagenomes from Israel (PRJNA828396), South Korea (PRJNA945504), China (PRJNA429990), the United States (PRJNA400072 and PRJNA389280), the Netherlands (PRJNA400072), and Spain (PRJEB1220), as well as two independent cohorts in the validation set from the United States (PRJNA398089) and Hong Kong (PRJNA1086048). For datasets containing an excessively large number of samples, a subset of samples was randomly selected to ensure computational feasibility. Sample selection was performed independently of any biological or environmental characteristics.

### Metagenome preprocessing and identification of viral contigs

Raw metagenomic reads from all cohorts were processed using a unified bioinformatics pipeline. Adapter sequences and PhiX174 spike-in reads were removed from the paired-end datasets using BBDuk (BBMap package; https://sourceforge.net/projects/bbmap/) with the following parameters: *ktrim=r*, *k=21*, *mink=11*, *hdist=1*, *qtrim=rl*, *trimq=18*, and *minlen=50*. Reads mapping to the human reference genome (hg38) were identified and removed using Kraken2^72^. High-quality, non-host reads were then assembled *de novo* with MEGAHIT v1.1.4^73^, using *k*-mer sizes ranging from 29 to 149 in increments of 10. Contigs shorter than 5 kbp were discarded prior to downstream analyses.

Putative viral contigs were identified using GeNomad^74^ with default parameters, and sequences classified as proviruses were excluded from subsequent analyses. Predicted viral sequences were then clustered into vOTUs using Vclust^75^. Pairwise average nucleotide identity (ANI) was calculated across all contigs, and clustering was performed with the Leiden algorithm (*--algorithm leiden*) using thresholds of 95% ANI and 85% query coverage (*--ani 0.95 --qcov 0.85*). The quality of each vOTU was assessed using CheckV v1.0.3^76^. Viral lifestyle classification was performed using PhaBOX v2.1.9^77^. Putative viral hosts were predicted with iPHoP v1.3.3^78^ with the iPHoP_db_Aug23_rw reference database (GTDB R214).

### Viral community analysis

To quantify viral abundance across samples, metagenomic reads were recruited against individual vOTUs. For computational efficiency, the vOTU catalog was partitioned into 50 batches (approximately 1,300 sequences per batch). Read mapping for each batch was performed using CoverM^79^ with a minimum read identity threshold of 97% (*--min-read-percent-identity 97*). The *--include-secondary* flag was enabled to retain secondary alignments, thereby capturing reads that map to shared genomic regions among closely related viruses. vOTUs were considered present in a sample when the covered fraction was higher than 0.5. Samples with fewer than one million mapped reads were excluded from subsequent analyses to ensure sufficient sequencing depth for quantitative comparisons. Viral abundances were then calculated as reads per kilobase per million mapped reads (RPKM). Bray–Curtis dissimilarities were computed using the vegdist function in the R package vegan (v.2.7-1). Community-level variation in virome composition was visualized using NMDS via the metaMDS function. Statistical significance of compositional differences among groups was assessed using PERMANOVA implemented in the adonis2 function of the vegan package, with group assignment specified as the explanatory variable.

### Identification of disease-associated viral signatures

To identify viral features associated with disease status, multivariable association testing was performed using MaAsLin2. Relative abundances of vOTUs were modeled against disease state as a fixed effect within a linear model (analysis_method = ‘LM’). Prior to model fitting, features were standardized (standardize = TRUE) to ensure comparability across samples. Statistical significance was determined at a 5% false discovery rate using the Benjamini–Hochberg correction. Model coefficients were interpreted as effect sizes, representing both the magnitude and direction of association, with positive coefficients indicating enrichment in patients and negative coefficients indicating enrichment in controls. Viral features with FDR-adjusted *p*-value < 0.05 were considered significant and retained for downstream analyses.

### Development of ML**-**based diagnostic models

The discovery dataset was randomly partitioned into a training set (80%) and an internal validation set (20%). Because the validation set was designed to evaluate performance across the two major geographic strata (Asian vs Western), all samples from the Israeli cohort were retained in the training set. A random forest classifier was built on the training set using the R package ranger (v.0.17.0), implemented through the machine-learning workflow provided by the R package caret (v.7.0-1). Class labels were encoded as a binary factor, with UC designated as the positive class. To mitigate class imbalance, synthetic minority oversampling (sampling = ‘smote’) was applied within repeated five-fold cross-validation (repeats = 3). Model training employed the Hellinger split rule with 1,500 trees, while *mtry* (20–40) and *min.node.size* (40–110) were tuned to assess parameter sensitivity and identify optimal model configurations. The optimal probability threshold for disease classification was determined by maximizing Youden’s index (Y = sensitivity + specificity − 1).

The best-performing model was applied to the validation subset and subsequently to independent datasets comprising cohorts from the United States and Hong Kong. Predictive performance was assessed using AUC, sensitivity, and specificity. AUC values were computed and visualized using the R package pROC (v.1.18.5).

### Gene prediction and functional annotation

Open reading frames (ORFs) were predicted from vOTUs, followed by sequence homology-based functional annotation using Pharokka^80^ with default parameters. For structure-based annotation, ORFs longer than 2.5 kb were excluded, as structural prediction accuracy decreases substantially for large proteins. The remaining ORFs were analyzed with Phold^40^, which integrates ProstT5-derived 3Di tokenization^41^ with Foldseek^42^ structural alignment to infer protein functions. Functional assignments were derived by querying multiple reference databases, including PHROG^81^ (viral protein families), VFDB^82^ (virulence factors), CARD^83^ (antibiotic resistance genes), DefenseFinder^84^ (prokaryotic defense systems), and NetFlax^85^ (toxin–antitoxin modules). Functional enrichment of VFGs and ARGs was assessed by Fisher’s exact test implemented in base R. Additional functional annotation of genes within two representative phage genomes (Fig. 6) was performed using HHpred^86^ against the PDB_mmCIF70_25_May database.

### Reporting summary

Further information on research design is available in the Nature Portfolio Reporting Summary linked to this article.

## Data availability

The raw metagenomic data have been deposited in the NCBI under the BioProject accession PRJNA1371390. All viral contigs and vOTUs, together with predicted gene annotations and corresponding protein sequences, are available on Figshare at https://figshare.com/account/articles/30900317. Source data are provided with this paper. Due to privacy and ethical restrictions, associated metadata cannot be made publicly available. However, metadata may be shared for bona fide research purposes upon approval from both our research institute and the requesting institution’s ethics committees. Requests for metadata access should be directed to the corresponding author, Jang-Cheon Cho.

## Code availability

The code and scripts used for data analyses are available via GitHub at https://github.com/HongjaePark-bio/gut_virome_ulcerative_colitis.

## AUTHOR CONTRIBUTIONS

J-C.C, I.K, H.K, and D-W.L conceptualized and designed the study. H.J, J.N, and Y.L contributed to sample collection and processing. H.P contributed to data interpretation, visualization, and writing the original manuscript draft. All authors reviewed and approved the manuscript.

## Competing interests

The authors declare no competing interests.

**Correspondence and requests for materials** should be addressed to Jang-Cheon Cho.

## Extended Data Figures

**Extended Data Fig. 1.**
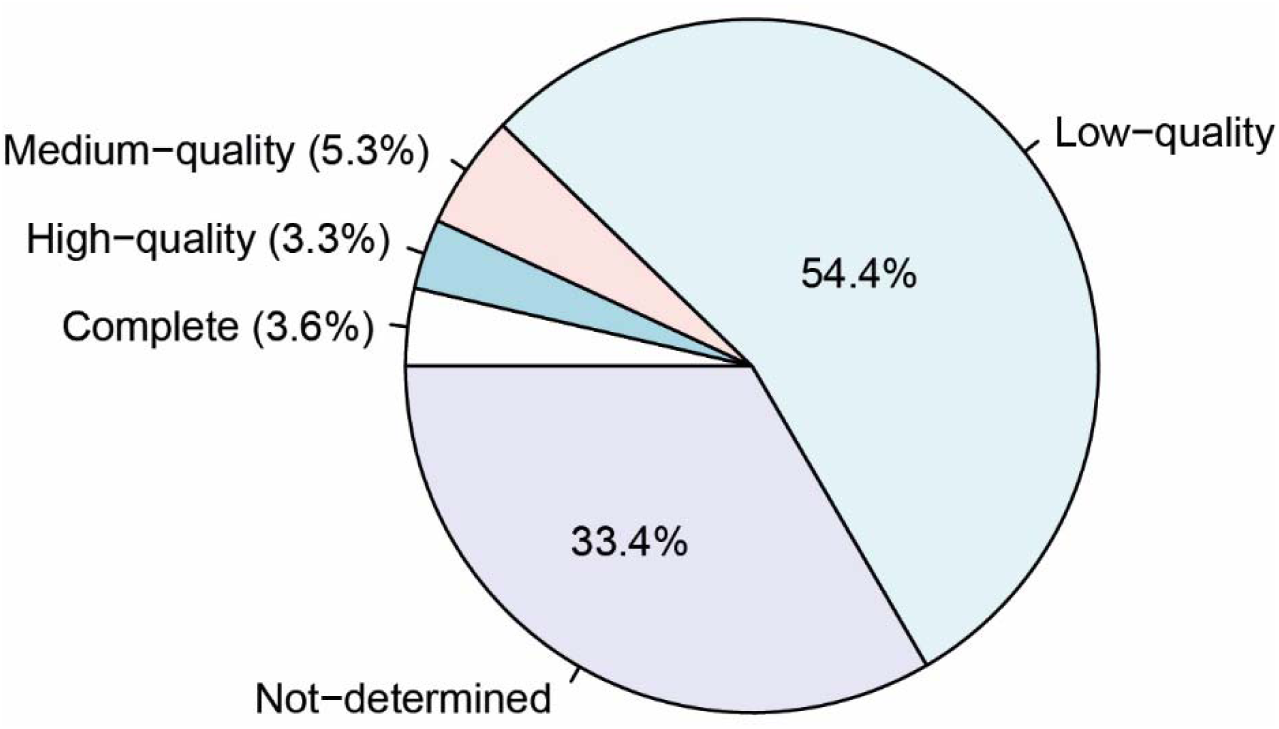
CheckV-based quality assessment of vOTUs. The 68,271 vOTUs were classified into five quality categories based on CheckV estimates: complete (100% completeness), high-quality (>90%), medium-quality (50–90%), low-quality (<50%), and not-determined (insufficient information for completeness estimation). Percentages indicate the proportion of vOTUs in each category.

**Extended Data Fig. 2.**
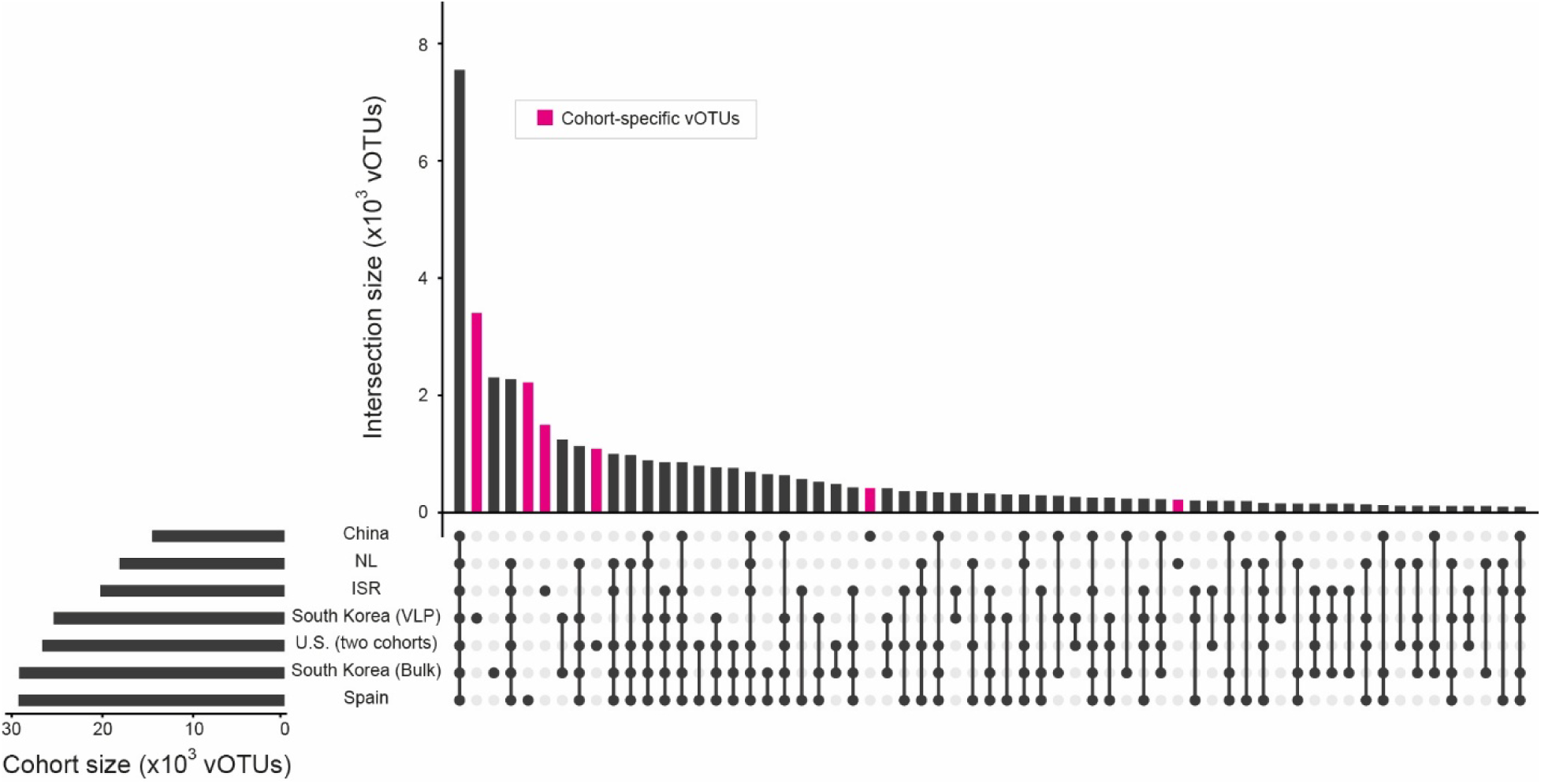
Cohort-specific and shared vOTUs identified in the discovery dataset. UpSet plot shows the presence–absence patterns of vOTUs across individual cohorts. Vertical bars indicate the number of vOTUs shared among the cohort combinations shown below, whereas horizontal bars represent the total number of vOTUs detected in each cohort. Cohort-specific vOTUs are highlighted in magenta. The two cohorts from the United States were combined for this analysis.

**Extended Data Fig. 3.**
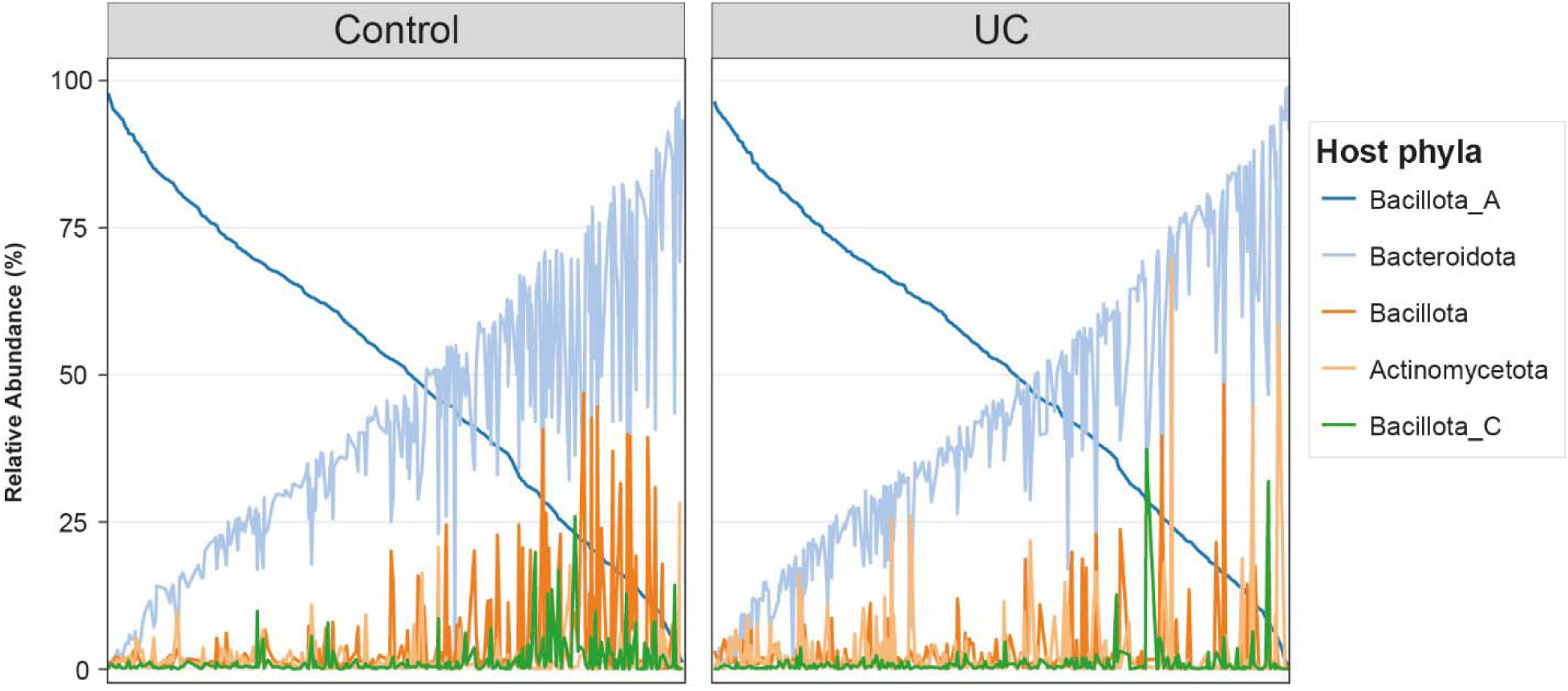
Relative abundances of the five most abundant viral groups, classified by predicted bacterial host phyla, in healthy controls and UC patients. Samples are grouped by disease status and ordered according to the relative abundance of phages predicted to infect *Bacillota_A*.

**Extended Data Fig. 4.**
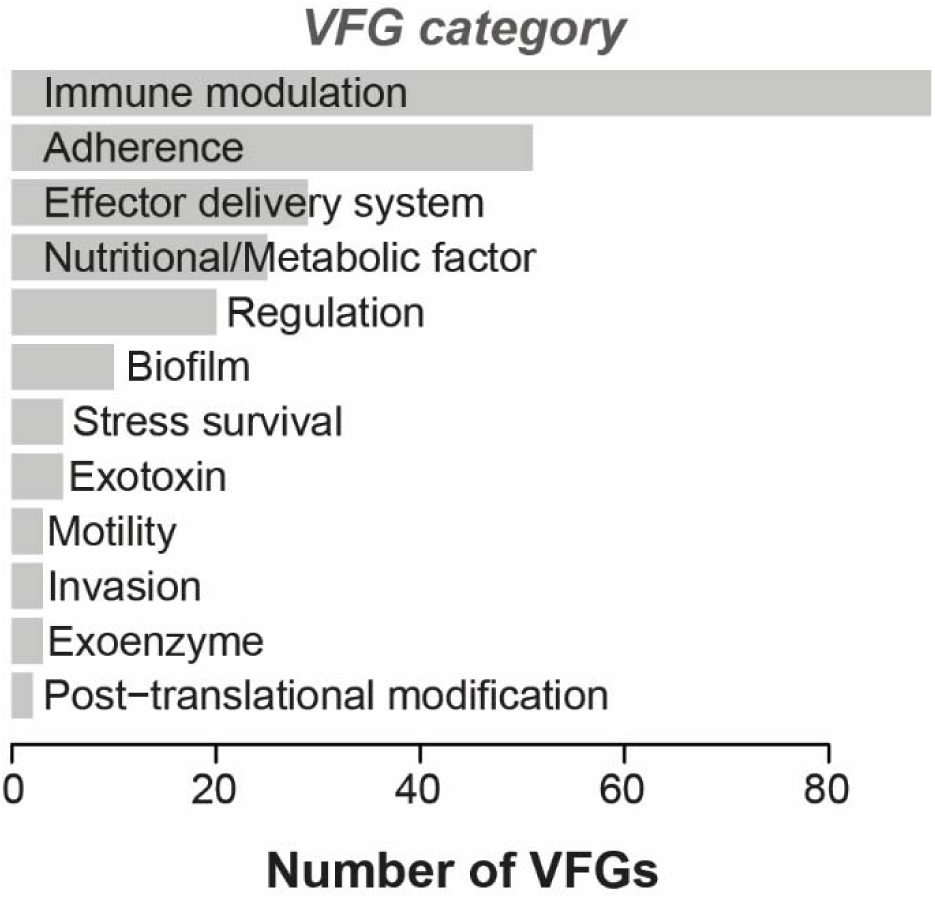
Distribution of VFGs in UC-depleted phages across major functional categories. Bar plot shows the number of VFGs assigned to each functional category in phages depleted in UC.

**Extended Data Fig. 5.**
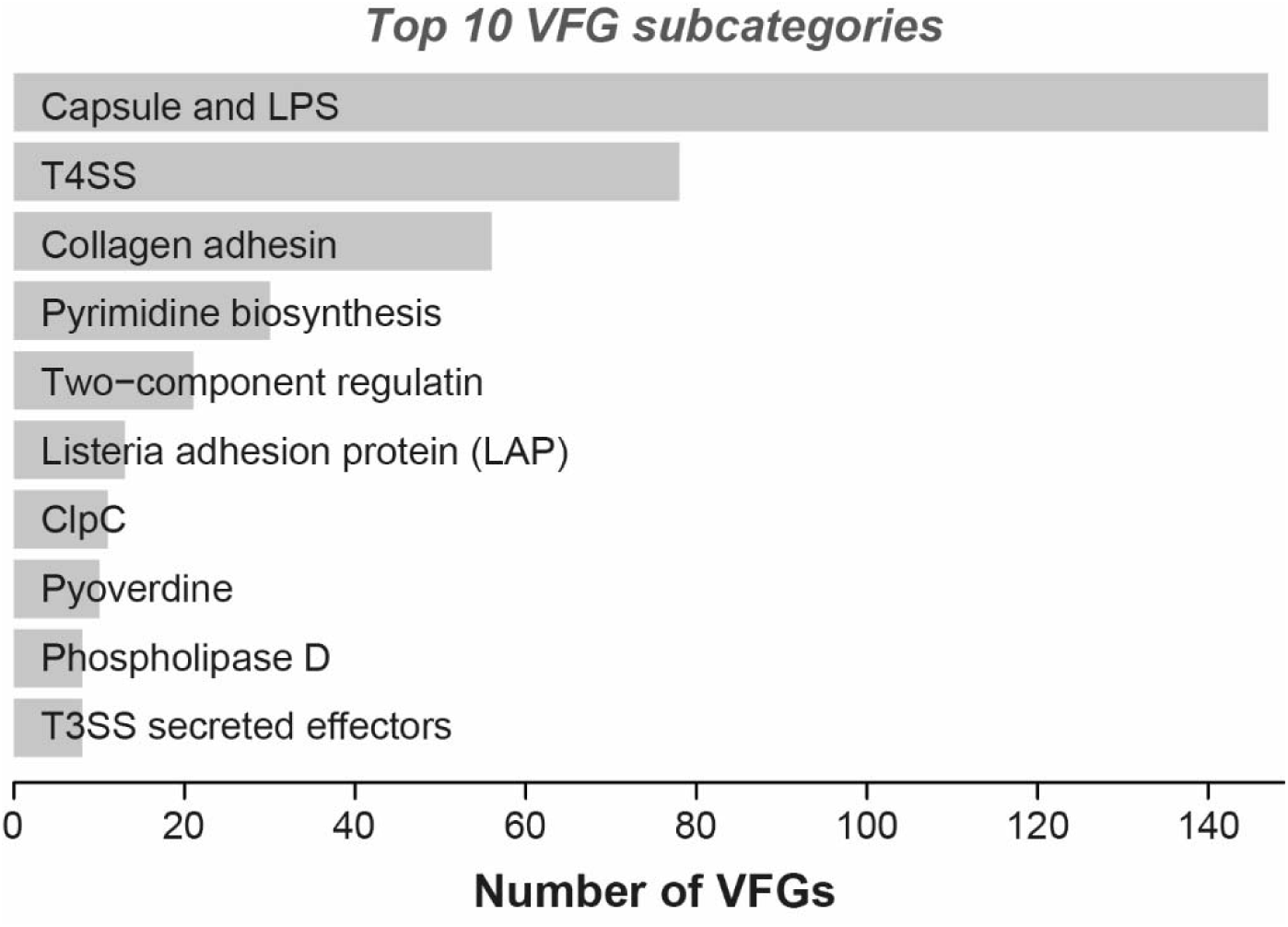
Distribution of VFGs in UC-enriched phages across functional subcategories. Bar plot shows the number of VFGs assigned to each functional subcategory in phages enriched in UC.

**Extended Data Fig. 6.**
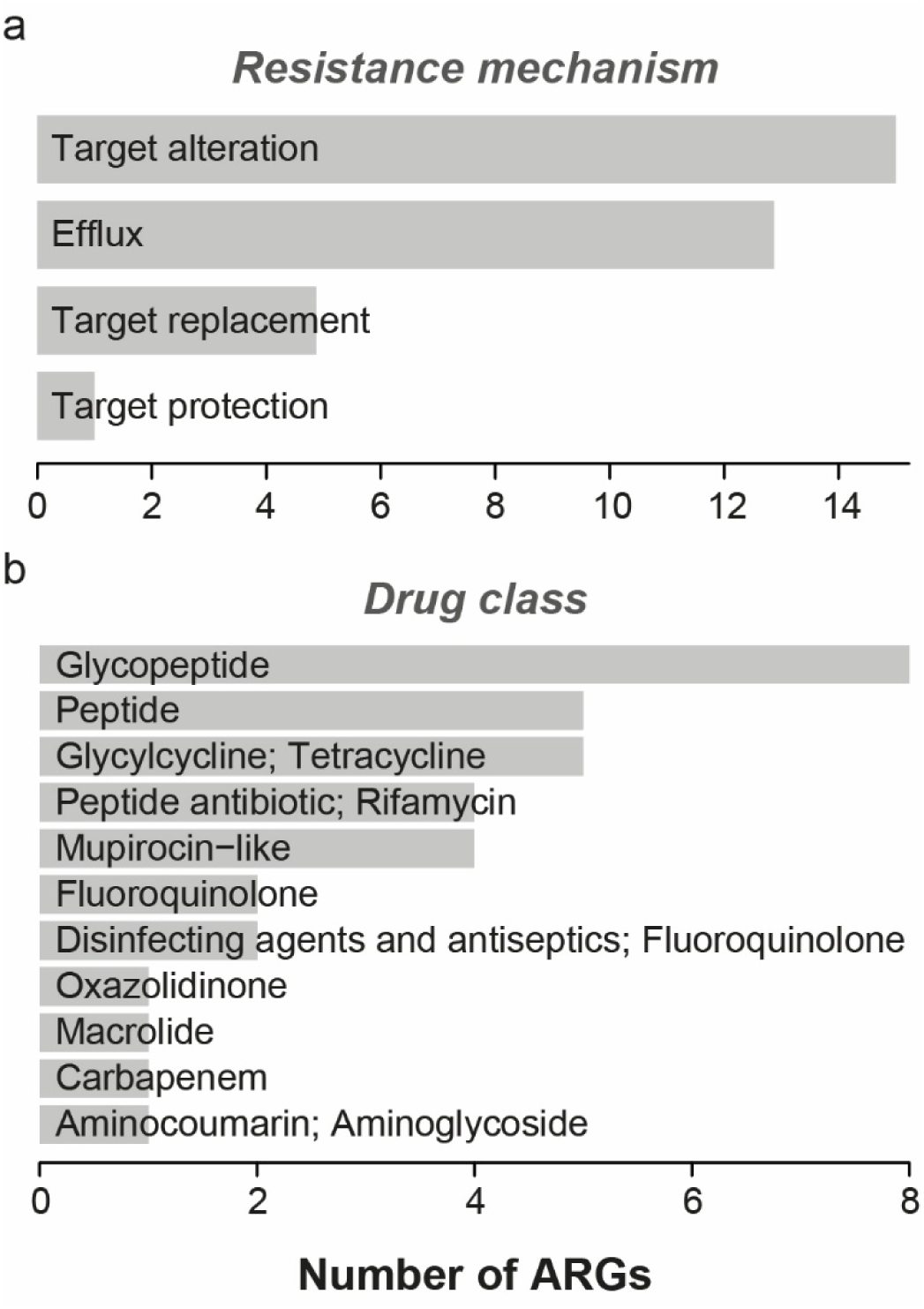
Distribution of ARGs in UC-depleted phages across resistance mechanisms and drug classes. Bar plots show the number of ARGs assigned to each resistance mechanism (a) and drug class (b) in phages depleted in UC.

**Extended Data Fig. 7.**
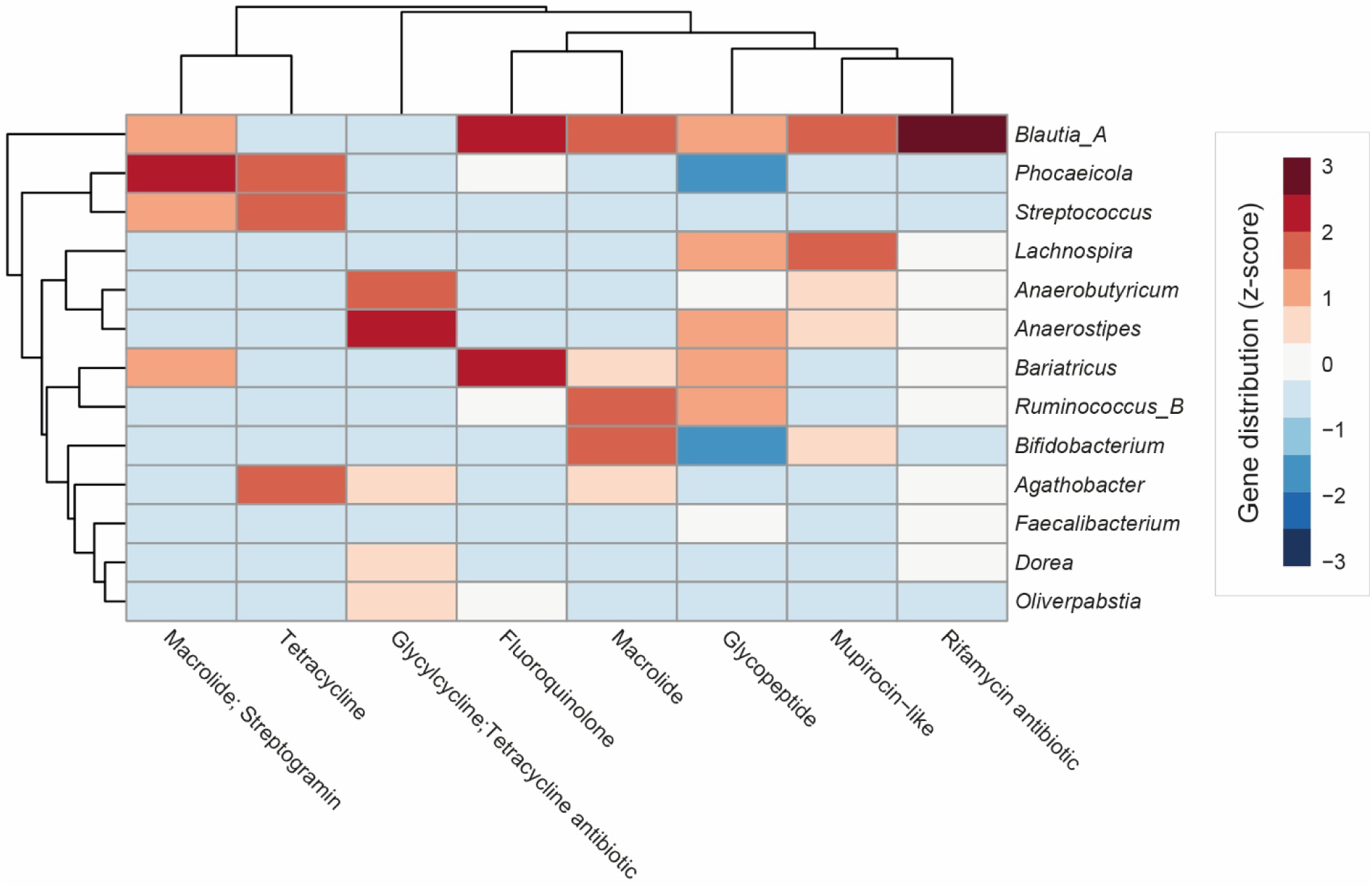
Distribution of ARGs in UC-enriched phages across predicted host bacterial genera. Heatmap shows the relative distribution (z-score) of ARGs, classified by drug class, across predicted host bacterial genera in phages enriched in UC.

